# Combining Dependent P-values with an Empirical Adaptation of Brown’s Method

**DOI:** 10.1101/029637

**Authors:** William Poole, David L. Gibbs, Ilya Shmulevich, Brady Bernard, Theo Knijnenburg

## Abstract

**Motivation:** Combining P-values from multiple statistical tests is a common exercise in bioinformatics. However, this procedure is non-trivial for dependent P-values. Here we discuss an empirical adaptation of Brown’s Method (an extension of Fisher’s Method) for combining dependent P-values which is appropriate for the correlated data sets found in high-throughput biological experiments.

**Results:** We show that Fisher’s Method is biased when used on dependent sets of P-values with both simulated data and gene expression data from The Cancer Genome Atlas (TCGA). When applied on the same data sets, the Empirical Brown’s Method provides a better null distribution and a more conservative result.

**Availability:** The Empirical Brown’s Method is available in Python, R, and MATLAB and can be obtained from https://github.com/IlyaLab/CombiningDependentPvaluesUsingEBM.1

## 1 Introduction

In order to integrate the large and diverse datasets found in systems biology, it is common to combine P-values from multiple statistical tests. The earliest method to combine independent P-values is seen in the work of [Fisher, 1948]. [Brown, 1975] extended Fisher’s Method to the case where P-values are assumed to be drawn from a multivariate normal distribution with a known covariance matrix. [Kost and McDermott, 2002] further extended Brown’s Method analytically for unknown covariance matrices. Additional methods for combining P-values have been developed for specific purposes, e.g. combining differently weighted P-values [Whitlock, 2005], combining P-values across multiple heterogeneous data sources [Aerts et al., 2006], and restricting analysis to the tail of the P-value distribution [Zaykin et al., 2002].

Of these methods, Brown’s most simply combines equally weighted dependent P-values. However, Brown’s reliance on multidimensional numerical integration makes it impractical for use on large data sets due to the computational resources needed to run the algorithm. Instead of using numerical integration, our adaptation of Brown’s Method uses the empirical cumulative distribution function derived from the data making our method dramatically more efficient and suitable for large omics data.

## 2 Methods

### 2.1 Fisher’s and Brown’s Methods

Let there be k P-values, denoted *P_i_,* generated from *k* statistical tests based on *k* normally distributed random variables. Fisher showed that for independent P-values, the statistic 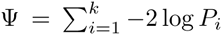 follows a chi squared distribution with 2*k* degrees of freedom, 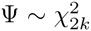. Brown extended Fisher’s Method to the dependent case by using a re-scaled χ^2^ distribution.

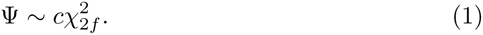

The constants *f* and *c* represent a re-scaled number of degrees of freedom and a scale factor which is the ratio between the degrees of freedom of Fisher’s and Brown’s methods. Brown calculated these constants by equating the first two moments of Ψ and 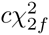 resulting in

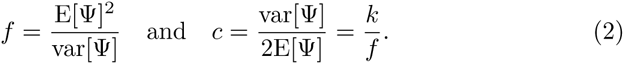

Furthermore, Brown showed that the expected value and variance of Ψ can be calculated directly via numerical integration to find the covariance, respectively;

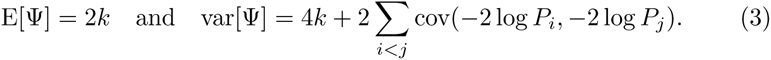

Numerical integration is needed to find the covariance term [Kost and McDermott, 2002]. The combined P-value is then given by *P*_combined_ = 1.0 − Ф_2_*_f_*(*ψ*/*c*) where 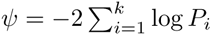 and Ф_2_*_f_* is the cumulative distribution function of 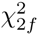.

### 2.2 Empirical Brown’s Method

Our contribution is to calculate the covariance in Eq. 3 empirically. In practice, each individual P-value, *P_i_*, will be computed via a statistical test between a variable and a vector of samples from the raw data, 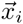. We define the transformed sample vector 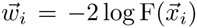 where 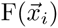 denotes the empirical cumulative distribution function calculated from the sample 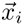. A more detailed explanation can be found in **SI1**. As a result, the covariance can also be computed empirically

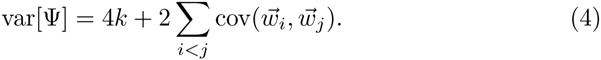

Line plot of histogram counts of P-values from Fisher’s Method applied on simulated null data with varying degrees of covariance as represented by *a*. The histogram was created by binning the P-values in 20 bins of size 0.05 from 0 to 1. **b)** Similar to **a** but for P-values derived with the Empirical Brown’s Method. **c)** A scatter plot comparing pathway association P-values for the gene *EGFR* in the GBM dataset from TCGA. Each circle represents one pathway. The radius is proportional to c, which reflects the intra-pathway correlation.

### 2.3 Generating Simulated Null Data

We compared our method to Fisher’s Method on generated random data. We generated these data by combining 20 P-values from the Pearson correlations between a sample of independent normal random variables with mean 0 and variance 1 (*n* = 200) and a sample (*n* = 200) of data generated from a 20-dimensional multivariate normal distribution centered around 0 with covariance matrix Σ; *σ_ii_* = 1 and *σ_i≠j_* = *a,* i.e. diagonal elements of 1 and off-diagonal elements of *a.* We calculated 100,000 combined P-values for each value of *a ∈* {0.0,0.25, 0.5,0.75}. We note that at least 100 data points are needed for accurate convergence of this method (**SI2**).

### 2.4 Combining P-values based on TCGA expression data

Fisher’s and the Empirical Brown’s Methods were compared on the highly correlated gene expression data of glioblastomas (GBM) from TCGA [Brennan et al., 2013]. Specifically, we derived combined P-values by associating the expression levels of the gene *EGFR* with the expression levels of the genes in the curated cancer signaling pathways as defined by the Pathway Interaction Database (PID) [Schaefer et al., 2009]. First, we computed P-values from pairwise Pearson correlations between *EGFR* and all genes in a pathway. (If *EGFR* was a member of the pathway, the correlation between *EGFR* and itself was excluded.) Then, we combined these P-values for each of the pathways using both Fisher’s and the Empirical Brown’s Methods. For more details see **SI3**.

## 3 Results and Discussion

### 3.1 Empirical Brown’s Method Conservative on Null Data

Combined P-values from randomly generated data should follow a uniform distribution. With Fisher’s Method, the number of extremely low and extremely high P-values are inflated as the intra-correlation of the normally distributed data set is increased (**Fig. 1a**). The inflation of low P-values results in a high number of false positives even for modest coupling in the covariance matrix. With the Empirical Brown’s Method, the distribution of P-values is slightly inflated in the middle of the interval [0,1] and deflated towards the low and especially the high values (**Fig. 1b**). This suggests that our method is a conservative estimate.

**Figure 1:**
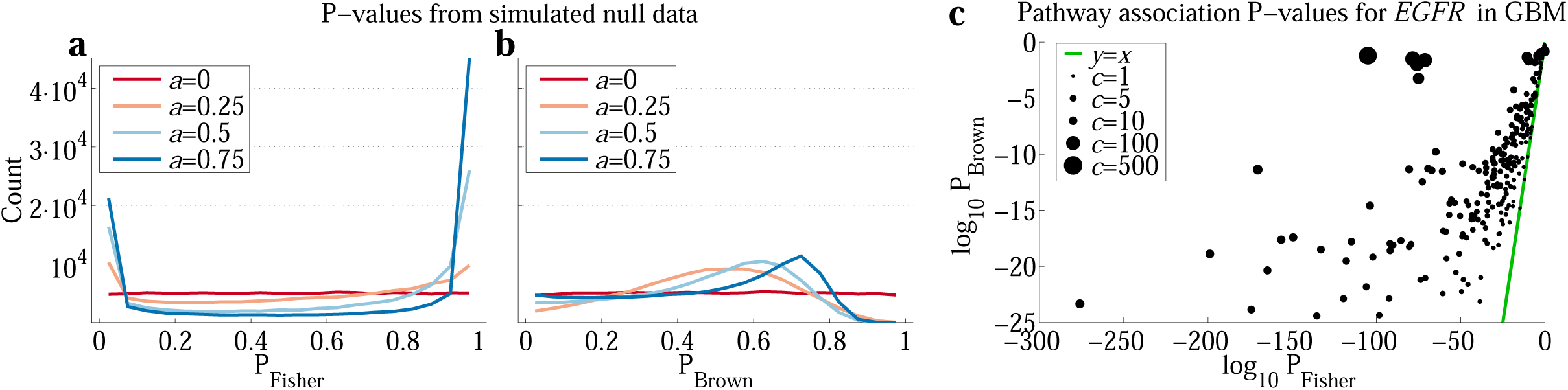
*P-values from simulated data and TCGA data using the Empirical Brown’s and Fisher’s Methods* a)

### 3.2 Our Method Corrects Fisher’s Bias on TCGA Data

As an example of combining dependent P-values generated from real intra-correlated data, we compared Fisher’s and the Empirical Brown’s Methods on associations between signaling pathways and *EGFR* using TCGA GBM gene expression data. *EGFR* is frequently amplified, mutated and overexpressed in GBM and is known to play an important functional role [Brennan et al., 2013]. It is therefore unsurprising that we observed many statistically significant associations between *EGFR* and the signaling pathways (**Fig. 1c** and **SI4**). However, Fisher’s Method produced much lower P-values, especially for the pathways with a high degree of intra-correlation, as quantified with the scale factor *c* (Eq. 2). This is a clear indication that Fisher’s Method produces spuriously low P-values when applied to correlated data. We also noted that Fisher’s Method produced very similar sets of significant pathways when correlated against a variety of genes other than *EGFR* (**SI4**). We interpret this as further evidence to suggest that Fisher’s Method is highly sensitive to the internal correlation structure of the data and detects significant associations in highly correlated sets of P-values regardless of the actual association. As seen in **Fig. 1c** and **SI4**, the Empirical Brown’s Method overcomes these biases.

## 4 Conclusion

On generated and real data, we show that the Empirical Brown’s Method overcomes biases in Fisher’s Method with regards to the internal correlation structure of the data used to generate the P-values. We believe that our implementations and evaluation of the method will provide a valuable tool for the bioinformatics community to combine P-values generated from statistically interdependent data sets.

## 6 Supplement SI1 Mathematical Explanation

Here we give a more detailed mathematical explanation of the Empirical Brown’s Method. We begin by explaining Brown’s method [Brown, 1975] in more detail largely following [Kost and McDermott, 2002]. Consider *k* normally distributed random variables with means 0 and covariance matrix Σ,

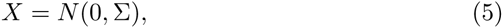
where *N*(0, Σ) is an *k*-dimensional normal distribution. P-values can be derived from *X* with with a cumulative distribution function,

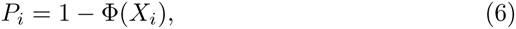
where *P_i_* denotes the *i^th^* P-value, *X_i_* denotes the *i^th^* component of *X*, and Φ denotes the cumulative distribution function. Note that this follows because the marginals of a multivariate normal distribution do not depend on the covariance. We now consider the distribution of

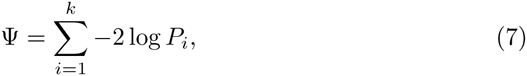
which we assume is proportional to a *χ*^2^ distribution with *2f* degrees of freedom, 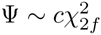. Brown showed that

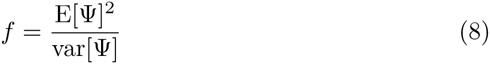
and

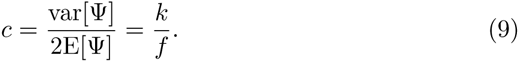

Assuming a *χ*^2^ distribution, E[Ψ] = 2*k*. Furthermore, define a new random variable *W_i_* = −2 log *P_i_* = −2 log (1 − Ф(*X_i_*)). Brown showed that,

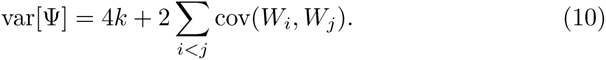

This expression can be evaluated for each *i* and *j* via numerical integration, where

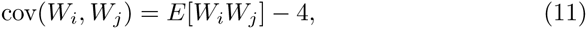

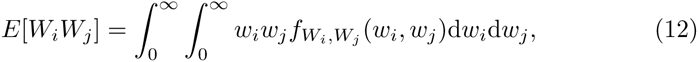
and 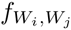 denotes the joint distribution between *W_i_.* and *W_j_*. Computationally, the challenge occurs when a large number of P-values are being combined - the number of numerical integrations scales with the square of the number of P-values being combined. Although this problem is parallelizable, it can still be computationally cumbersome for large data sets. Initial tests with numerical integration (not shown) in Python revealed that combining roughly 2000 P-values would take days with a single workstation, the bottleneck being these pairwise integration steps. Instead, we took an empirical approach and attempted to approximate cov(*W_i_, W_j_*) directly from the data. With this approach the computation only took hours on a workstation. Let 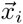 be a sample drawn from *X_i_*. We can approximate a sample, 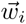, from *W_i_* by transforming the raw data using the empirical right-sided cumulative distribution function *F*,

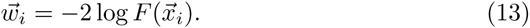

The covariance between two variables *W_i_* and *W_j_* can then estimated from the raw data using the well known definition of covariance,

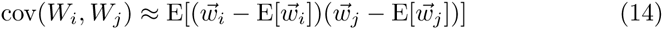

Due to the efficiency of many existing implementations for calculating the empirical cumulative distribution and the covariance, this method is practical for use on large data sets.

## 7 Supplement SI2 Null Distribution Generation and Convergence as a Function of Sample Size

To generate a single combined P-value, we combined *k* = 20 P values computed between *k* normally distributed random variables and *k*-dimensional vectors sampled (*n* = 200) from a dependent *k* dimensional normal distribution with covariance matrix given by

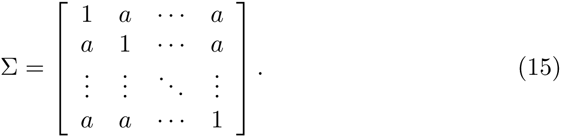

We computed 100,000 P-values for each value of *a* we considered; these data are shown in **Fig. 1a** in the main text.

Additionally, we found that given *n >* 100 samples per data vector 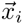, our implementation produces relatively little variation for the values of *f* and *c*. These tests were done by generating sample data from numerous example covariance matrices with known actual degrees of freedom (**Supp. Fig. 1**).

## 8 Supplement SI3 TCGA Data Analysis

### 8.1 Pathway Definitions

The NCI Nature Curated Pathway Interaction Database (PID) database consists of an ontology of pathways. In order to avoid additional unnecessary multiple testing, we restricted our set of pathways to leaf pathways in the pathway ontology tree [Schaefer et al., 2009].

**Supp. Fig. 1:**
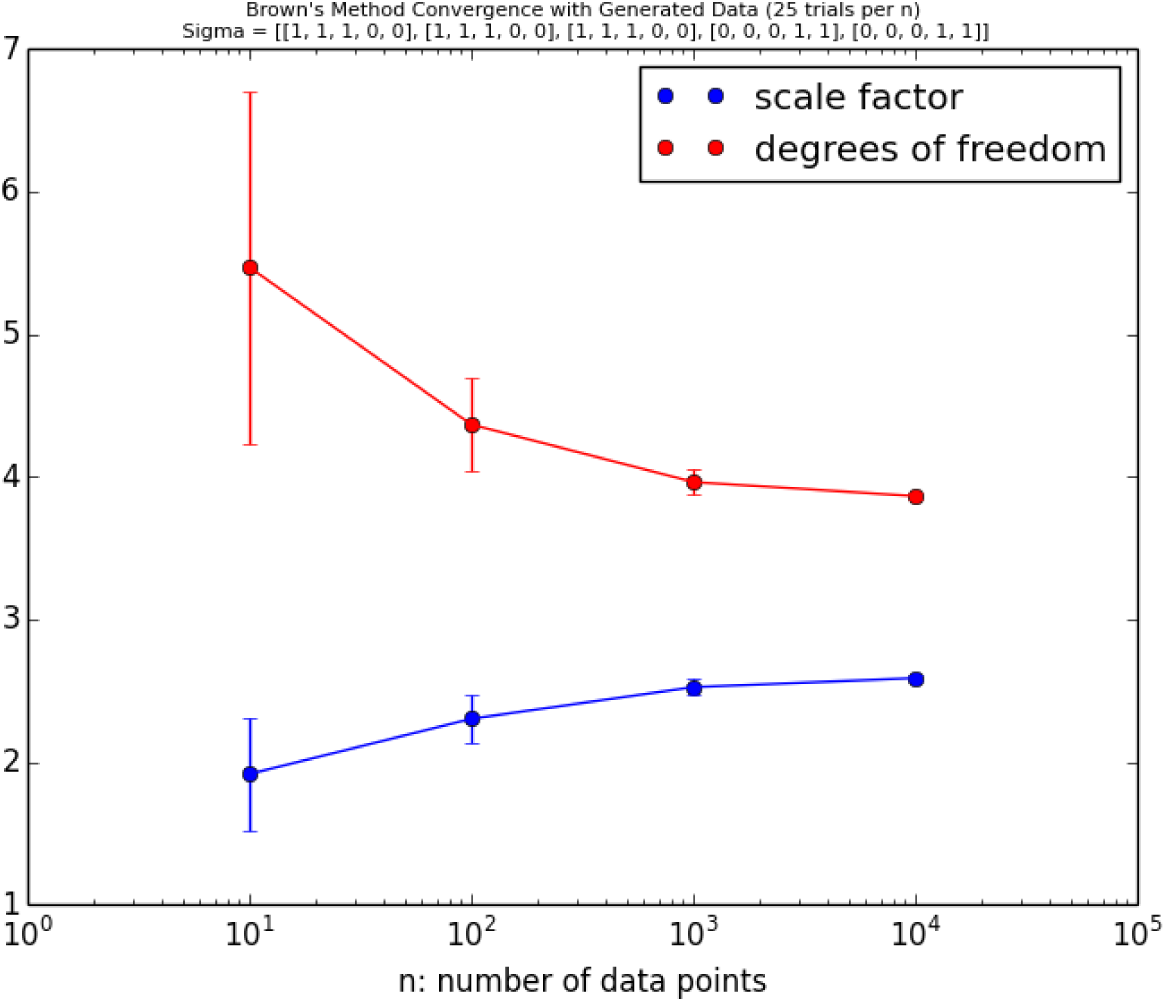
Convergence of Empirical Brown’s Method as a function of the sample size *n* when calculating *c* and *f*. A 5 by 5 covariance matrix with 2 degrees of freedom 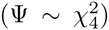 was used in this example. Error bars show standard deviation across 25 different trials.

**Supp. Fig. 2:**
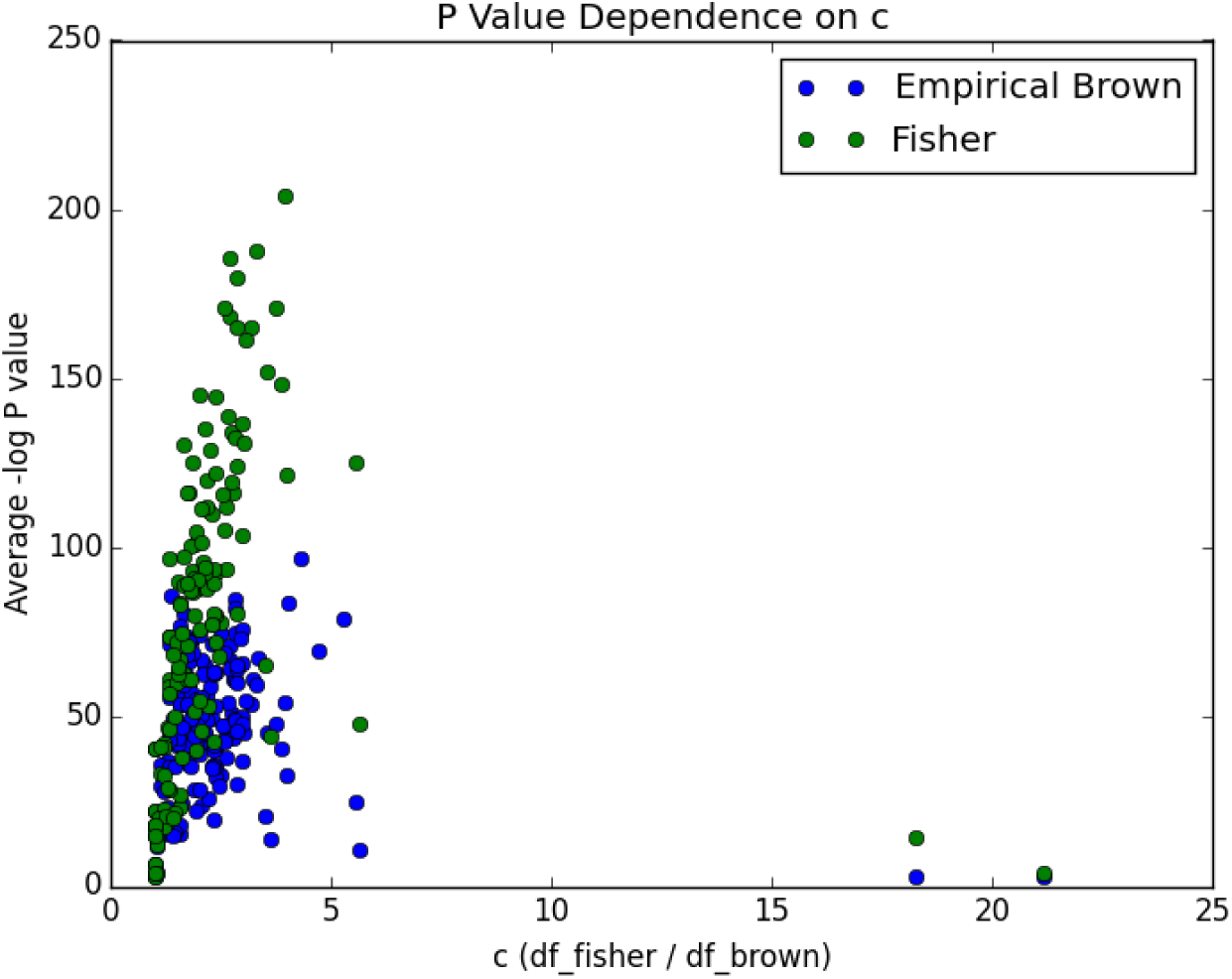
Each dot represents the average -log P-value for a single pathway. Notice the strong dependence on *c* when combining P-values with Fisher’s method. This dependence nearly vanishes when using our method.

### 8.2 Pathway Associations

In order to evaluate the Empirical Brown and Fisher’s Methods on TCGA data, we first computed a pairwise correlation matrix between pairs of genes based on their expression levels in glioblastoma samples (GBM) from TCGA [Brennan et al., 2013]. We restricted ourselves to genes in PID. We precomputed the entire covariance matrix between transformed GEXP data for increased efficiency. We then combined P-values on the pathway level. Let *C_P_* be the set of pairwise correlation P-values between *g*_0_ and the genes in each pathway, *C_P_* = {*p*_cor_(*g*_0_, *g_i_*); *g_i_* ∈ *P,g*_0_ ≠ *g_i_*}, where *P* is the set of genes in a pathway and *p*_cor_ denotes the P-value from the Pearson correlation computed via a two-tailed test of the t-distribution. The P-values in each set *C_P_* were combined using Fisher’s method and the Empirical Brown’s method. This analysis was done for each of the genes in PID. In the main text we have only reported on pathway associations with *EGFR.*

### 8.3 Note on Transforming Combined P-values to Q-values

A common way to correct for multiple testing other than the Bonferroni correction is by transforming the P-value to a Q-value (or false discovery rate). The two most common approaches to compute Q-values are Benjamini-Hochberg’s approach [Benjamini and Hochberg, 1995] and Storey’s approach [Storey, 2002]. Importantly, we noted that Storey’s approach to compute Q-values (as implemented in the R and Python packages [Storey et al., 2015]) is not directly appropriate for significance testing in this case due to complications in estimating the null hypothesis distribution. Specifically, the implementation of Storey’s approach assumes a particular distribution on the P-values and estimates parameters to fit this distribution. The P-value distributions encountered in the gene expression data were problematic in terms of estimating these parameters, leading to nonsensical results. Benjamini-Hochberg’s non-parametric approach does not suffer from this limitation.

**Supp. Fig. 3:**
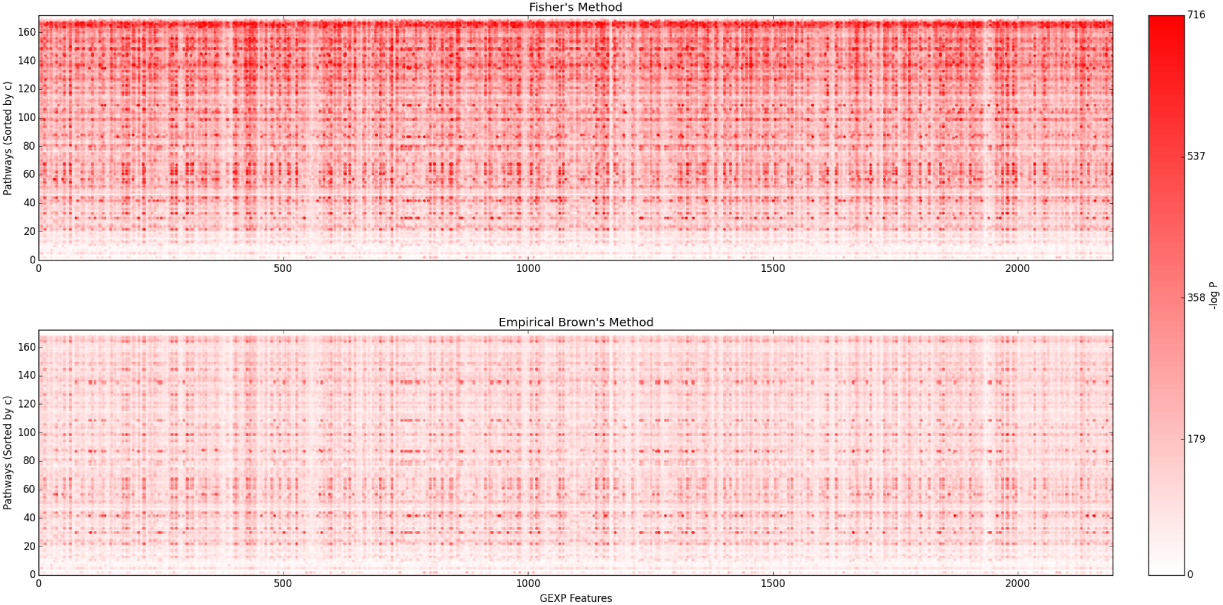
Each red dot represents a significant GEXP feature pathway association. Pathways are sorted by their intra-pathway correlation, quantified by *c* from low to high

## 9 Supplement SI4 P-Value Dependence on *c*

Fisher’s method shows a strong bias towards more intra-correlated pathways. The relative degrees of freedom between Fisher’s and the Empirical Brown’s Method provides a good measure of the correlation within a pathway, because this value quantifies the change percentage of degrees of freedom (in terms of variables), which are statistically redundant due to correlations with other variables. We can see this effect in two ways. First, **Supp. Fig. 2** shows that the average -log P-value in Fisher’s method across all genes is strongly correlated with *c*. The Empirical Brown’s Method, on the other hand, does not show this statistical dependence. Second, the top occurring significant pathways are frequently shared across genes. **Supp. Fig. 3** shows genes (x-axis) and pathways (y-axis) with significant associations in red. The pathways are sorted by c from low to high. Notice that pathways with higher values of c tend to be more red when using Fisher’s Method. Our method diminishes this bias.

## 10 Supplement SI5 Software Implementation

Our implementation of the Empirical Brown’s Method in Python uses the scipy, numpy, and statsmodels libraries. This implementation is efficient and is easily applicable to large scale genomics data. Let there be k data vectors denoted 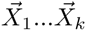 each with *n* samples. Our function takes for input a matrix of these data vectors and a vector of *k* P-values, denoted *P*_1_ *… P*_k_, to be combined. The works as follows:

- Transform the data to its normal coordinates - mean of 0 and unit variance 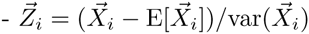.
- Calculate the empirical cumulative distribution function (F) over the data using the statsmodel package.
- Approximate the −2 log cumulative distribution vector, for each data vector; 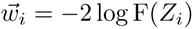.
- For each pair of indices (*i, j*) calculate the covariance cov 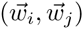.
- Sum covariances to calculate var[Ψ], *f* and *c*.
- Calculate the combined statistic 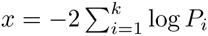.
- Compute a meta P-value using Brown’s re-scaled distribution: 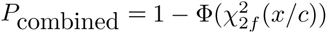, where Ф denotes the cumulative distribution function.

Additionally, for flexibility of use each component of our code can be called individually. This allows for the covariance matrix to be pre-computed and Brown’s method to be applied on arbitrary subsets of the data (which is how we carried out our TCGA analysis).

An efficient implementation is also available in R and Matlab. See https://github.com/IlyaLab/CombiningDependentPvaluesUsingEBM.

## References

Aerts, S. et al. (2006). Gene prioritization through genomic data fusion. Nature biotechnology, 24(5):537–544.

Benjamini, Y. and Hochberg, Y. (1995). Controlling the false discovery rate: a practical and powerful approach to multiple testing. Journal of the Royal Statistical Society. Series B (Methodological), pages 289–300.

Brennan, C. W. et al. (2013). The somatic genomic landscape of glioblastoma. Cell, 155(2):462–477.

Brown, M. B. (1975). 400: A method for combining non-independent, one-sided tests of significance. Biometrics, pages 987–992.

Fisher, R. A. (1948). Answer to question 14 on combining independent tests of significance. The American Statistician, 2(30).

Kost, J. T. and McDermott, M. P. (2002). Combining dependent p-values. Statistics & Probability Letters, 60(2):183–190.

Schaefer, C. F. et al. (2009). Pid: the pathway interaction database. Nucleic acids research, 37(suppl 1):D674–D679.

Storey, J., Bass, A., Dabney, A., and Robinson, D. (2015). Package qvalue.

Storey, J. D. (2002). A direct approach to false discovery rates. Journal of the Royal Statistical Society: Series B (Statistical Methodology), 64(3):479–498.

Whitlock, M. (2005). Combining probability from independent tests: the weighted z-method is superior to fisher’s approach. Journal of evolutionary biology, 18(5):1368–1373.

Zaykin, D. V. et al. (2002). Truncated product method for combining p-values. Genetic epidemiology, 22(2):170–185.

